# Cross-Platform Omics Pipeline for Integrated Transcriptomics

**DOI:** 10.1101/2025.06.11.659031

**Authors:** Lianchong Gao, Huangying Le, Xin Zou, Jie Hao

## Abstract

The fragmentation of single-cell RNA-seq (scRNA-seq) and spatial transcriptomics (ST) tools across Python/R ecosystems, coupled with escalating computational demands for million-cell datasets, creates critical barriers to reproducible analysis. To overcome this, we present CrosPIT (Cross-Platform Omics Pipeline for Integrated Transcriptomics), a cloud-native framework that unifies cross-language analysis in Google Colab with versioned storage on Google Drive. Pre-configured environments enable rapid switching between CPU/GPU/R runtimes, eliminating local hardware constraints. To demonstrate biological utility, we embedded several modules, such as a module for precision detection of malignant epithelial cells in colorectal cancer.

CrosPIT thus delivers a scalable, language-agnostic solution that standardizes workflows through native AnnData/Seurat object interoperability, accelerates integrative omics research, and democratizes access to high-performance transcriptomic analysis.

## Introduction

Rapid advances in single-cell RNA sequencing (scRNA-seq) and spatial transcriptomics (ST) have provided unprecedented resolution for analyzing cellular heterogeneity^1-5^, but the resulting complex analysis workflows pose major challenges for researchers^6,7^. Mainstream analysis tools are scattered across different programming ecosystems such as Scanpy^8^ in Python and Seurat^9^ in R, and cross-platform collaboration often requires manual data format conversion and rewriting of analysis scripts, which leads to fragmented workflows and threatens reproducibility. As datasets with millions of cells become increasingly common, computationally intensive tasks such as dimensionality reduction and clustering place high demands on local hardware resources; many researchers are constrained by insufficient computing power or lack of high-performance GPUs. Finally, compatibility issues among data formats (e.g., AnnData, RDS, Loom) and disorganized management of intermediate files further exacerbate the risk of information loss during multi-stage analyses.

To address these challenges, we developed **Cr**oss-Platform **O**mic**s P**ipeline for **I**ntegrated **T**ranscriptomics (CrosPIT), an end-to-end single-cell analysis pipeline built on the Google Colab cloud platform with structured storage on Google Drive. This pipeline allows users to analyze million-cell datasets via online servers without requiring local high-performance computing resources, while also supporting deployment on local computing clusters with dedicated hardware. The core design principle of the framework is cross-language workflow integration, which optimizes the transcriptomic analysis paradigm.

Specifically, CrosPIT offers the following advantages. First, the Colab cloud platform provides a unified runtime environment, and we supply standardized environment configuration scripts or pre-configured environments, enabling users to deploy their analysis environment conveniently and efficiently. Second, by using Google Drive as structured storage and leveraging Colab’s ability to flexibly switch between programming environments and hardware platforms at runtime, CrosPIT enables complex analyses across languages and platforms with elastic resource scheduling. Third, the framework integrates the tools of each analysis stage into a standardized workflow, and provides complete reproducible example code and detailed tutorials. Fourth, its modular design ensures extensibility: each step outputs canonical Seurat or AnnData objects, allowing users to pass intermediate results directly to downstream algorithms or visualization tools without additional conversion, thereby facilitating rapid adoption of emerging methods.

### CrosPIT facilitates cross-language, cross-platform data analysis

CrosPIT is an end-to-end scRNA-seq and ST analysis pipeline built on Google Colab with structured storage on Google Drive, designed to enable users to perform data analysis across programming environments and hardware platforms (Fig. 1A). Leveraging Colab, CrosPIT supports switching between Python and R runtime environments, integrating both language ecosystems into a single workflow. Colab can flexibly allocate computing resources of different capacities to suit various hardware needs, and it provides data format conversion functions to facilitate data exchange between analysis tools (Fig. 1B, 1C). Additionally, CrosPIT uses Google Drive for structured management of intermediate files, and a unified naming convention and version control ensure traceability of the analysis process and reproducibility of results. Finally, relying on the stability of the Colab runtime environment, CrosPIT enables rapid cloud environment deployment via pre-configured environment files or unified setup scripts, minimizing the time required for initial configuration. Together, these features allow CrosPIT to flexibly conduct single-cell data analysis across different software ecosystems and hardware platforms, laying the groundwork for further applications in integrated analyses of scRNA-seq and ST data. For example, processing a large sample might involve the following workflow. First, raw single-cell data are stored on Google Drive. Next, a Colab runtime with 300 GB of RAM is connected, and the environment is configured rapidly. Cell Ranger^10^ preprocessing, CellBender^11^ background correction, and scVelo^12^ analysis are then performed sequentially to obtain the initial processed data files. The runtime is then switched to one with GPU support, and the environment is reconfigured quickly. A Scanpy pipeline accelerated by the rapid-singlecell package on GPU is used to perform data quality control, batch integration, dimensionality reduction, clustering, and annotation, yielding a single-cell reference dataset in h5ad format. Finally, the runtime is switched to an R environment with another quick setup, the h5ad file is converted to RDS format, and RCTD^13^ uses the single-cell reference data to deconvolute the ST dataset. In this workflow, switching between runtimes is easily accomplished through the Colab web interface.

**Figure 1.**
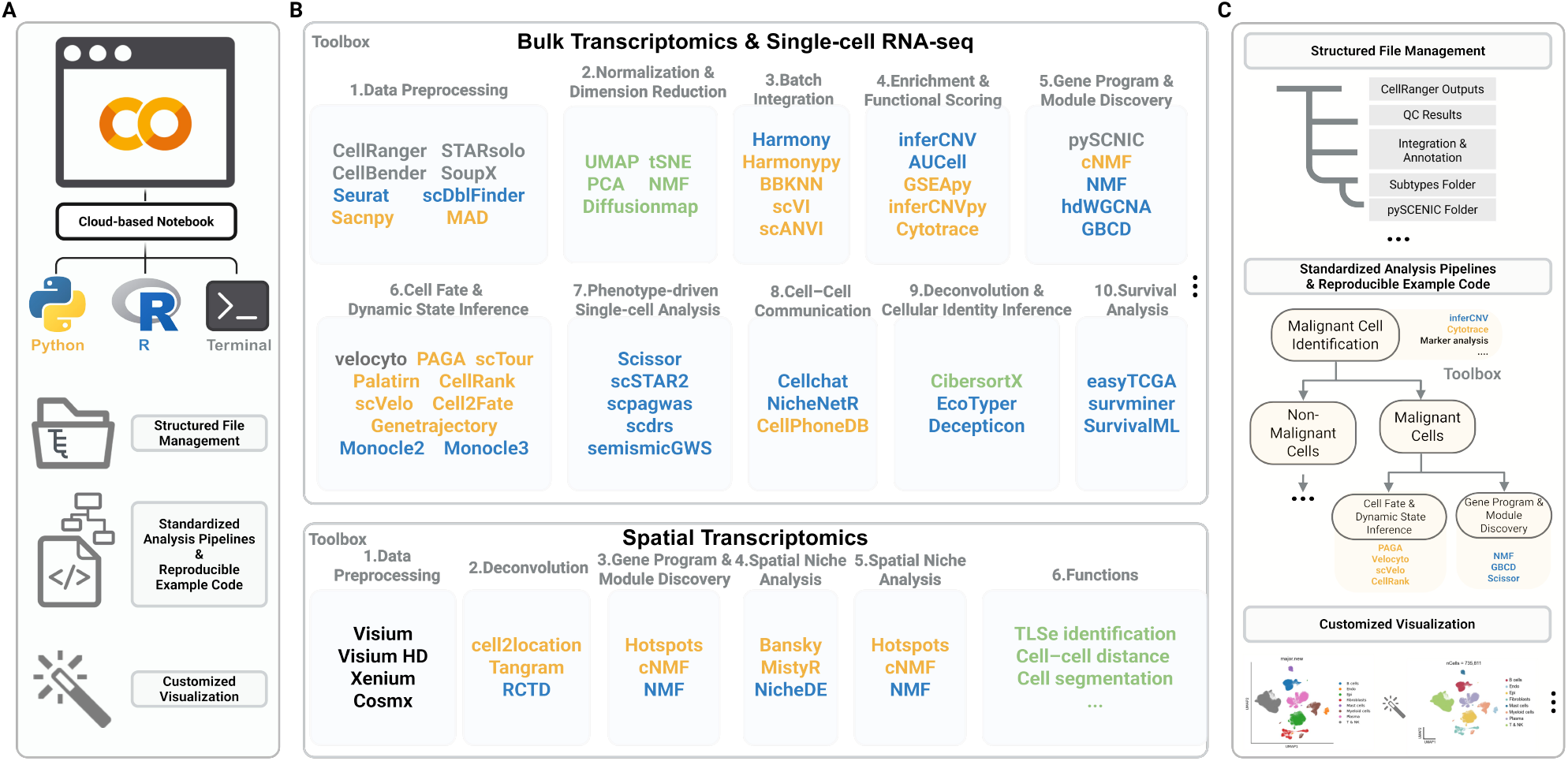
Architecture and key components of the CrosPIT pipeline. **A**. Cloud-based execution environment. CrosPIT runs in a Google Colab notebook that can switch seamlessly between Python and R kernels or a Unix-style terminal. Pre-configured environment files allow one-click deployment. **B**. Integrated multi-omics toolbox. CrosPIT bundles representative open-source tools for every major step of bulk transcriptomics, single-cell RNA-seq and spatial-omics analysis. Coloured tool names indicate the primary language ecosystem (orange = Python, blue = R, black = language-agnostic/CLI). **C**. Standardised file and workflow management. CrosPIT enforces a hierarchical directory structure and ships reproducible template notebooks. Example sub-pipelines (shown for malignant-cell identification) inherit this structure, feed intermediate objects directly into downstream modules and output publication-ready visualisations.

All important intermediate files are output to Google Drive so that they can be accessed by each runtime. Moreover, the environment configuration files are highly stable, enabling rapid environment setup.

Therefore, CrosPIT not only provides infrastructure for integrating multimodal data such as scRNA-seq and ST, but more importantly establishes an extensible cross-platform collaboration paradigm. In addition, emerging algorithms can be rapidly deployed using existing toolchains without reengineering the underlying data formats, since the pipeline is built on mainstream data structures such as AnnData (Python) and Seurat (R) objects.

### Automated malignant-cell identification in CrosPIT

Within CrosPIT, we implement an integrated pipeline for identifying malignant cells in colorectal cancer (CRC) using single-cell RNA-sequencing data^14,15^ (Fig. 2A). First, after integrated clustering of multiple samples with Seurat or Scanpy, epithelial clusters are extracted and patient-specific indices (Fig. 2B–D) are calculated for EPCAM- and KRT8-positive clusters. These indices include cluster purity (proportion of cells from the same patient), the adjusted Rand index (ARI), normalized mutual information (NMI) and the silhouette coefficient; clusters with high values indicate patient-specific malignant populations. Using epithelial cells from matched normal tissue or non-epithelial cells as references, CNV inference is then performed with inferCNV^16^ or inferCNVpy (|CNV| > 0.1) and CopyKAT^17^ to detect aneuploidy (Fig. 2B–D). Module scores are simultaneously computed for CRC lineage markers (EPCAM, KRT20), oncogenes (CEACAM5, KRAS) and stemness genes (PROM1/CD133); a malignancy score—tumour gene-set minus normal gene-set (> 0)—provides additional evidence. For genomically stable CRC cells, the CNV threshold is lowered and identification relies on characteristic transcriptional features (Fig. 2B–D).

**Figure 2.**
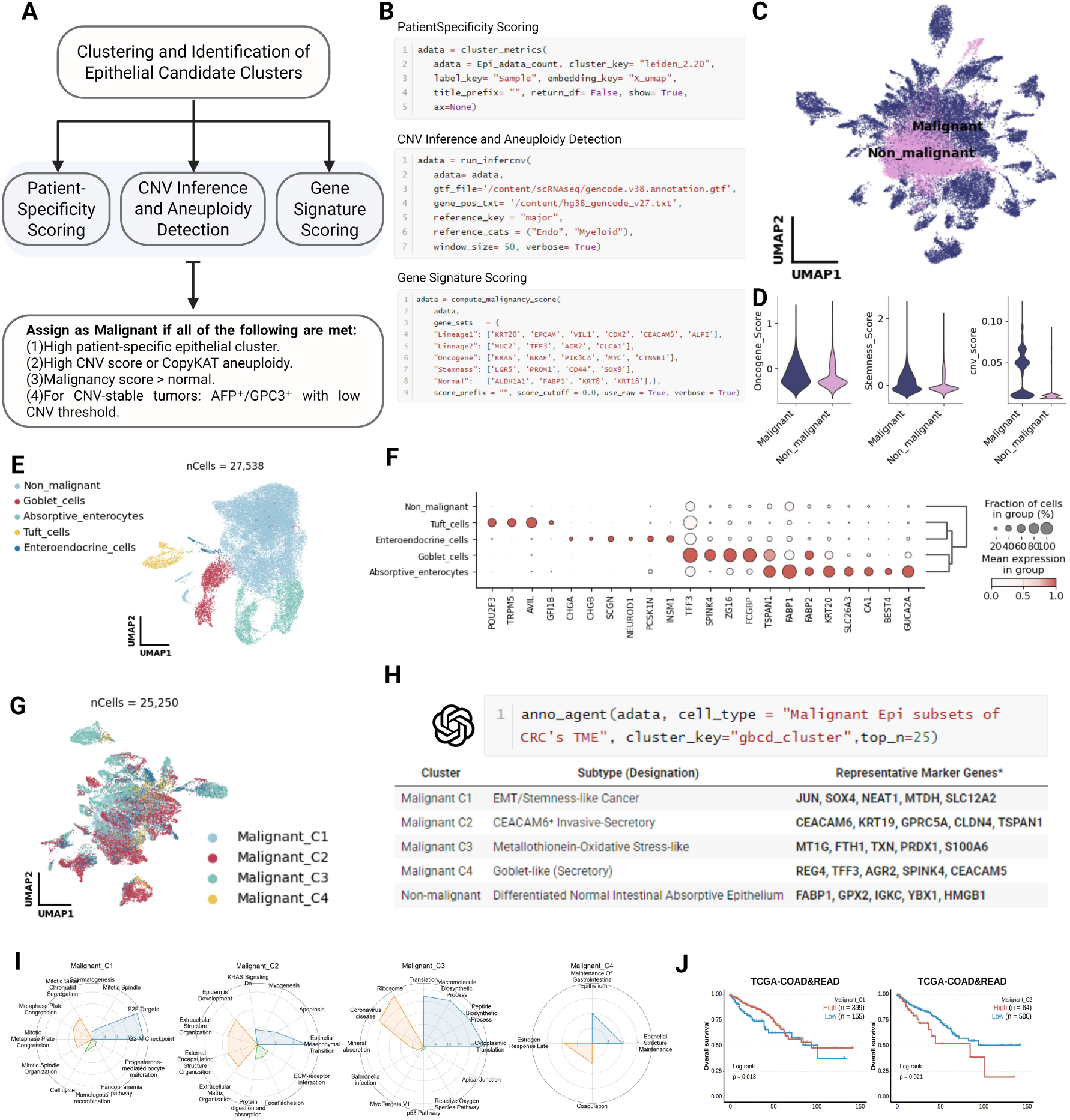
CrosPIT module for automated identification and characterisation of malignant epithelial cells, illustrated on a colorectal-cancer (CRC) dataset. **A**. Decision logic. Epithelial clusters are first scored for patient-specificity, copy-number variation (CNV)/aneuploidy and malignant gene-signature activity. A cell is labelled malignant only if it satisfies all four criteria shown in the lower box. **B**. Key code snippets. Example one-liners (Python) for computing patient-specificity metrics (top), running CNV inference with inferCNVpy (middle) and calculating malignancy-related module scores (bottom). **C**. Global UMAP. All epithelial cells coloured by the final malignant-versus-non-malignant assignment. **D**. Discriminatory scores. Violin plots for CNV score, malignancy score and stemness/onco-gene module scores, contrasting malignant and non-malignant cells. **E**. Non-malignant epithelial landscape. UMAP after removing malignant cells, revealing five differentiated intestinal lineages. **F**. Marker validation. Dot plot showing the expression of canonical lineage markers across non-malignant subtypes. **G**. Malignant sub-clustering. UMAP of 28 817 malignant cells re-clustered into four transcriptional programmes (Malignant C1–C4). **H**. Automated subtype annotation. Output from anno_agent, listing each malignant cluster, its inferred subtype and representative marker genes. **I**. Functional profiling. Radar charts summarising pathway or signature enrichments for each malignant cluster. **J**. Clinical relevance. Kaplan–Meier survival curves (TCGA-COAD&READ) stratified by high-versus low-expression of cluster-specific signatures, illustrating prognostic differences.

In the automated decision framework, a cell is classified as malignant only if it simultaneously satisfies: (1) membership in an epithelial cluster with a high patient-specificity index; (2) a CNV score above threshold or aneuploid status by CopyKAT; (3) a malignant gene-module score exceeding the normal-tissue score; and (4) for CNV-stable tumours, a relaxed CNV cut-off. CrosPIT then generates publication-ready visualisations—including UMAPs, CNV heat-maps and marker-gene bubble plots—and exposes key parameters such as the CNV threshold and clustering resolution for user adjustment. Preset marker-gene lists for ten solid-tumour types ensure immediate compatibility across cancers without re-engineering.

After malignant cells are removed, the residual epithelial compartment resolves into five differentiated intestinal lineages (Fig. 2E, F). The malignant fraction is re-clustered into four transcriptional programmes, Malignant C1–C4 (Fig. 2G), which are automatically annotated by the ‘anno_agent’ module with representative marker genes (Fig. 2H). Radar plots highlight pathway and signature enrichments for each malignant subtype (Fig. 2I), and Kaplan–Meier analyses using TCGA-COAD/READ demonstrate their distinct prognostic implications (Fig. 2J).

## Data and code availability

The scRNA-seq data for CRC are deposited under accession number GSE178341, GSE132465 and GSE144735. CrosPIT can be accessed at the following GitHub repository:https://github.com/LCGaoZzz/CrosPIT.git.

## Acknowledgements

The authors express their gratitude for the provision of data by databases such as TCGA and GEO.

